# EEG-Neurofeedback Targeting Gamma Oscillations at the Parieto-Occipital Region Reduces Pain Perception

**DOI:** 10.64898/2026.05.27.728092

**Authors:** Xiangyue Zhao, Hao Sun, Shiyu Wei, Haoqing Duan, Gan Huang, Jingwei Li, Hailu Wang, Xuejing Lu, Yanzhi Bi, Li Hu

## Abstract

Pain is closely associated with both *spontaneous* and stimulus-*evoked* gamma oscillations, which differ in their phenomenology and their relationship to pain perception. These associations make gamma oscillations a promising target for pain interventions, such as closed-loop real-time neurofeedback (NFB). While modulating stimulus-*evoked* gamma oscillations is technically challenging due to their concurrent occurrence with pain perception, targeting *spontaneous* gamma oscillations is both feasible and practical, as it could influence subsequent cortical nociceptive processing and reduce pain. To test this mechanistic hypothesis, we developed a novel NFB training protocol aimed at increasing *spontaneous* gamma oscillations in the parieto-occipital region to alleviate pain. Eight-eight healthy, right-handed subjects were randomly assigned to either an active or sham NFB group and completed three sessions of NFB training. During each session, subjects underwent EEG recording and received randomized laser stimulation while watching a video. After three training sessions, approximately 52% of subjects in the active group successfully increased the spectral power of *spontaneous* gamma oscillations in the parieto-occipital region. Crucially, the spectral power of *spontaneous* gamma oscillations was significantly and negatively correlated with subjective pain intensity after the third NFB session. Additionally, subjects who responded successfully to active NFB showed significant reductions in pain intensity, unpleasantness, and laser-evoked potentials compared to the sham group. Together, these findings establish a causal relationship between *spontaneous* gamma oscillations in the parieto-occipital region and pain perception, potentially offering a novel NFB-based therapeutic strategy for pain management.

## 1. Introduction

Pain is a major global health issue, affecting millions of individuals and leading to profound physical, emotional, and economic consequences^1–4^. While pharmacological treatments, particularly those involving opioids, are widely used to relieve pain, they often carry significant risks, including side effects and addiction, highlighting the urgent need for alternative pain management strategies^5^. Recent advances in understanding the neural mechanisms of pain and the rapid progress in neuromodulation techniques create opportunities to develop targeted, mechanism-based pain relief strategies, enabling more precise and personalized treatments^6–8^.

Neural oscillations are rhythmic neural activities in the central nervous system that underpin cognitive functions and link microscopic neuronal dynamics to macroscopic behavioral outputs^9–16^. Increasing evidence suggests a close relationship between pain perception and neural oscillations^9,17–20^, particularly gamma oscillations^21–29^. Specifically, pain is associated with both *spontaneous* and stimulus-*evoked* gamma oscillations, which differ in their phenomenology^30^, generation mechanisms^31^, and their relationship to pain perception^28^. Nociceptive-*evoked* gamma oscillations, originating in the superficial layers of the primary somatosensory cortex (S1) contralateral to nociceptive stimulation, reflect neural activity coupled with the spike firing of interneurons^32^. These oscillations can not only track subjective pain ratings within individuals but also predict inter-individual variations in pain sensitivity^29,33,34^. In contrast, spontaneous gamma oscillations, largely governed by inhibitory interneurons, are distributed across various brain regions^35^. Modulating *spontaneous* gamma oscillations in different brain areas can directly influence the associated behaviors^31^. Our previous studies have consistently demonstrated that *spontaneous* gamma oscillations in the parieto-occipital region prior to nociceptive stimulation significantly and inversely correlated with subsequent pain perception^28,36^. These findings suggest that *spontaneous* gamma oscillations could be used as a promising target for pain interventions, such as transcranial alternating current stimulation (tACS) and closed-loop real-time neurofeedback (NFB).

Notably, tACS has been used to target gamma oscillations at 50/80 Hz in the sensorimotor cortex, which is the neural origin of nociceptive-*evoked* gamma oscillations. However, no significant analgesic effects were observed^37^. This lack of efficacy may stem from difficulties in effectively modulating gamma oscillations during pain perception. In contrast, NFB provides real-time feedback on neural activity, enabling individuals to self-regulate their oscillations in a specific direction to influence behaviors^38,39^. This capacity for real-time, targeted modulation makes NFB particularly well-suited for investigating the causal role of *spontaneous* gamma oscillations in pain perception.

To explore the causal relationship between *spontaneous* gamma oscillations and pain, we implemented a three-session NFB training protocol to enhance *spontaneous* gamma oscillations in the parieto-occipital region. We hypothesize that repeated NFB training will enhance *spontaneous* gamma oscillations, subsequently reducing both pain perception and nociceptive-evoked brain responses. To rule out the possibility that modulation of *spontaneous* gamma oscillations during NFB training was driven by control of neck muscle activity, we simultaneously recorded electromyography (EMG) and electroencephalography (EEG) signals and assessed their correlations. This approach may also provide preliminary neuroscientific evidence for the development of novel pain modulation interventions.

## 2. Methods

### 2.1 Subjects

This study was approved by the Institutional Review Board of the Institute of Psychology, Chinese Academy of Sciences, and conducted in accordance with the Declaration of Helsinki. All subjects provided written informed consent and received financial compensation for their participation.

An *a priori* power analysis using G*Power software was conducted to determine the appropriate sample size for a mixed design (2 × 3 = 6 conditions). It yielded a sample size of n = 44 to detect a medium effect size of f = 0.25 at a standard error probability of α = 0.05 with a power of 0.95. To account for the anticipated presence of neurofeedback non-responders, a total of 88 healthy right-handed subjects were recruited for the two parts of the experiment (24 females; age = 22.55 ± 2.30 years, mean ± SD). Exclusion criteria included any psychiatric or mental disorders, chronic pain, substance abuse, irregular sleep patterns in the past week, or alcohol consumption within 24 hours prior to the experiment.

### 2.2 Experimental procedure

Subjects were randomly assigned to one of two groups and were blinded to their group allocation: an active group, which received real-time gamma feedback based on their own amplitudes, and a sham group, which received gamma feedback based on amplitudes of the preceding participant. The experiment consisted of two parts. In the first part, 44 subjects underwent the following procedures in sequence: determination of stimulus energy (∼10 min), pain assessment before NFB training (∼3 min), three sessions of NFB training (∼50 min), and pain assessment after NFB training (∼3 min, Figure 1A). In the second part, an additional 44 subjects were recruited. Besides completing the same experimental procedure, these subjects also underwent simultaneous posterior neck EMG recording during EEG acquisition to exclude the possibility that gamma activity over the parieto-occipital region reflected contamination from neck muscle activity. To assess task engagement and the credibility of the feedback manipulation, all subjects completed 10-point numerical rating scales (NRS), including measures of motivation before training, as well as effort and perceived feedback authenticity after training. The study protocol and planned analyses for the first part were retrospectively registered on the Open Science Framework (OSF). Prior to the initiation of the second part of the experiment, the OSF record was updated to provide a more complete and transparent description of the EMG recordings. The preregistration is available on OSF (10.17605/OSF.IO/2SRQA). All experimental procedures adhered to the standard protocol for neurofeedback experiments^40^ (Supplementary Materials).

**Figure 1.**
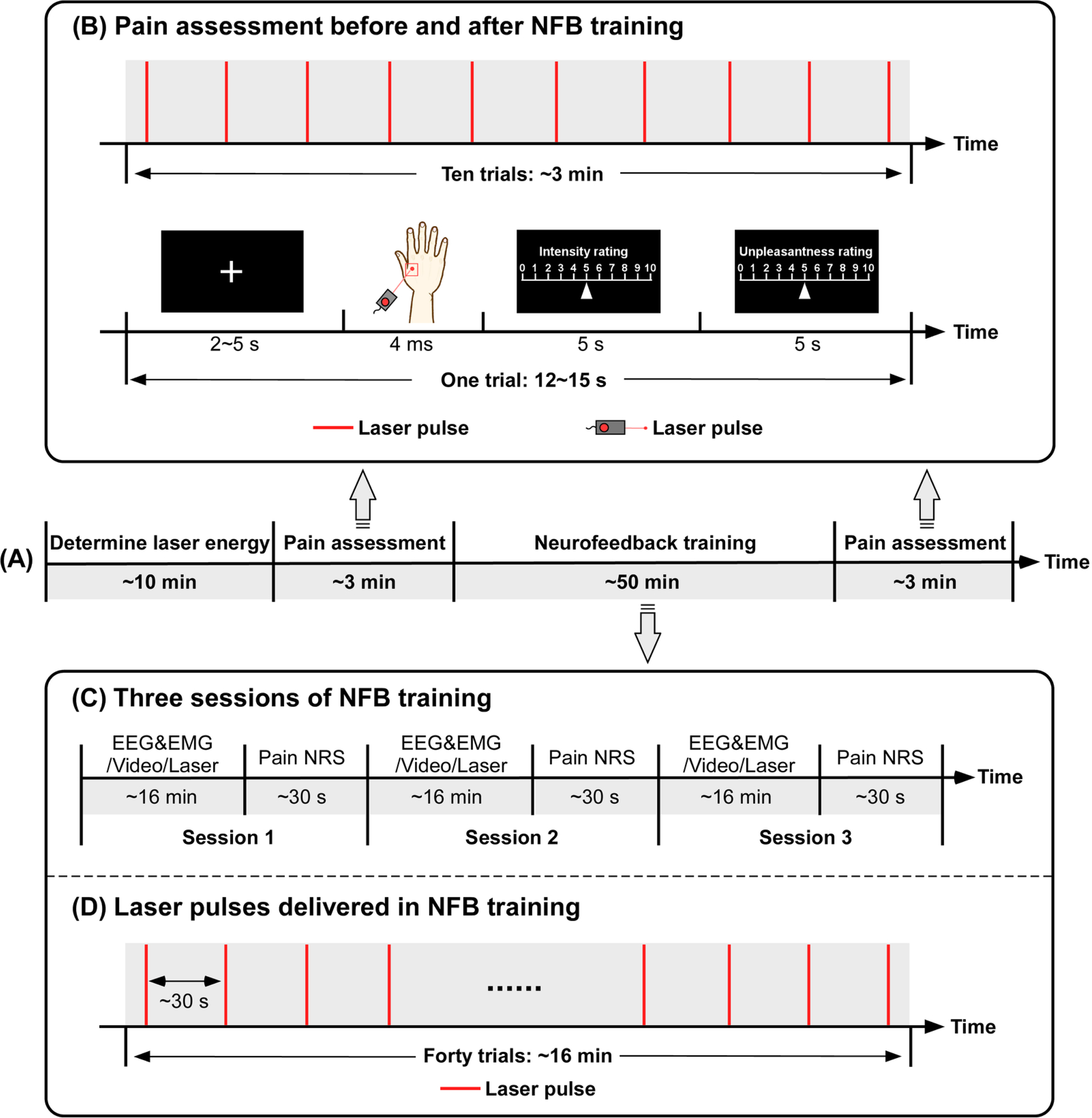
Experimental design. (A) Experimental procedure. All subjects completed a laser energy evaluation, pre-training pain assessment, NFB training, and post-training pain assessment. (B) Pain assessment before and after NFB training. Ten laser pulses were delivered to the dorsum of the right hand, with pain intensity and unpleasantness ratings recorded before and after NFB training for each participant. (C) Three sessions of NFB training. During each training session, EEG and EMG (in the second part of the experiment) data were recorded, while subjects watched a video and received nociceptive laser stimuli. After each session, subjects rated their pain intensity and unpleasantness. (D) Laser pulses delivered during NFB training. Forty laser stimuli were delivered to the dorsum of the participant’s right hand in each NFB training session.

#### 2.2.1 Nociceptive laser stimulation

Nociceptive stimuli were radiant heat pulses generated by an infrared neodymium yttrium aluminum perovskite (Nd:YAP) laser (Electronical Engineering, Italy). The laser emitted at a wavelength of 1.34 μm with a pulse duration of 4 ms^41^. The He-Ne laser pulses were transmitted through an optical fiber and focused by lenses to a spot with a diameter of ∼7 mm. Laser pulses were applied within a 5 × 5 cm² squared area on the dorsum of the subject’s right hand. The laser beam target was manually displaced by at least 1 cm in a random direction to prevent fatigue or sensitization of nociceptors. The ascending method of limits was adopted for each subject to determine the laser energy that could evoke a subjective pain rating of ∼6 on a NRS ranging from 0 (‘no pain’) to 10 (‘the worst pain imaginable’)^42,43^. This method was achieved by incrementally increasing the laser energy from 1.25 J in steps of 0.25 J until the target rating was obtained. This procedure was repeated three times, and the average energy that elicited a pain rating of ∼6 was used in subsequent experimental procedures.

#### 2.2.2 Pain assessments before and after NFB training

Ten laser pulses were delivered to the dorsum of the right hand to evaluate pain intensity and unpleasantness for each subject before and after the NFB training (Figure 1B). Each trial began with a 2-5 s presentation of a white cross centered on the screen, followed by the delivery of laser stimulus. After the stimulus, a visual cue was presented to instruct subjects to rate their perceived pain intensity on the 0-10 NRS within 5 s. Another visual cue then prompted them to rate their perceived unpleasantness on the same 0-10 NRS within 5 s. The duration of each trial varied randomly between 12 and 15 s, and each pain assessment session lasted ∼3 min in total.

#### 2.2.3 Online NFB training

The NFB training consisted of three sessions, each lasting ∼16 min. During each session, subjects were instructed to self-regulate their gamma oscillations while watching a video with real-time visual feedback (Figure 1C). In the meantime, forty laser stimuli were delivered at randomized intervals of about 30 s (Figure 1D) while recording EEG signals. Subjects were instructed to rate their perceived overall pain intensity and unpleasantness immediately after each training session. A schematic overview of the NFB training system for a single session is shown in Figure 2, which was composed of three main modules: (1) EEG and EMG recording, (2) gamma power calculation, and (3) visual feedback.

**Figure 2.**
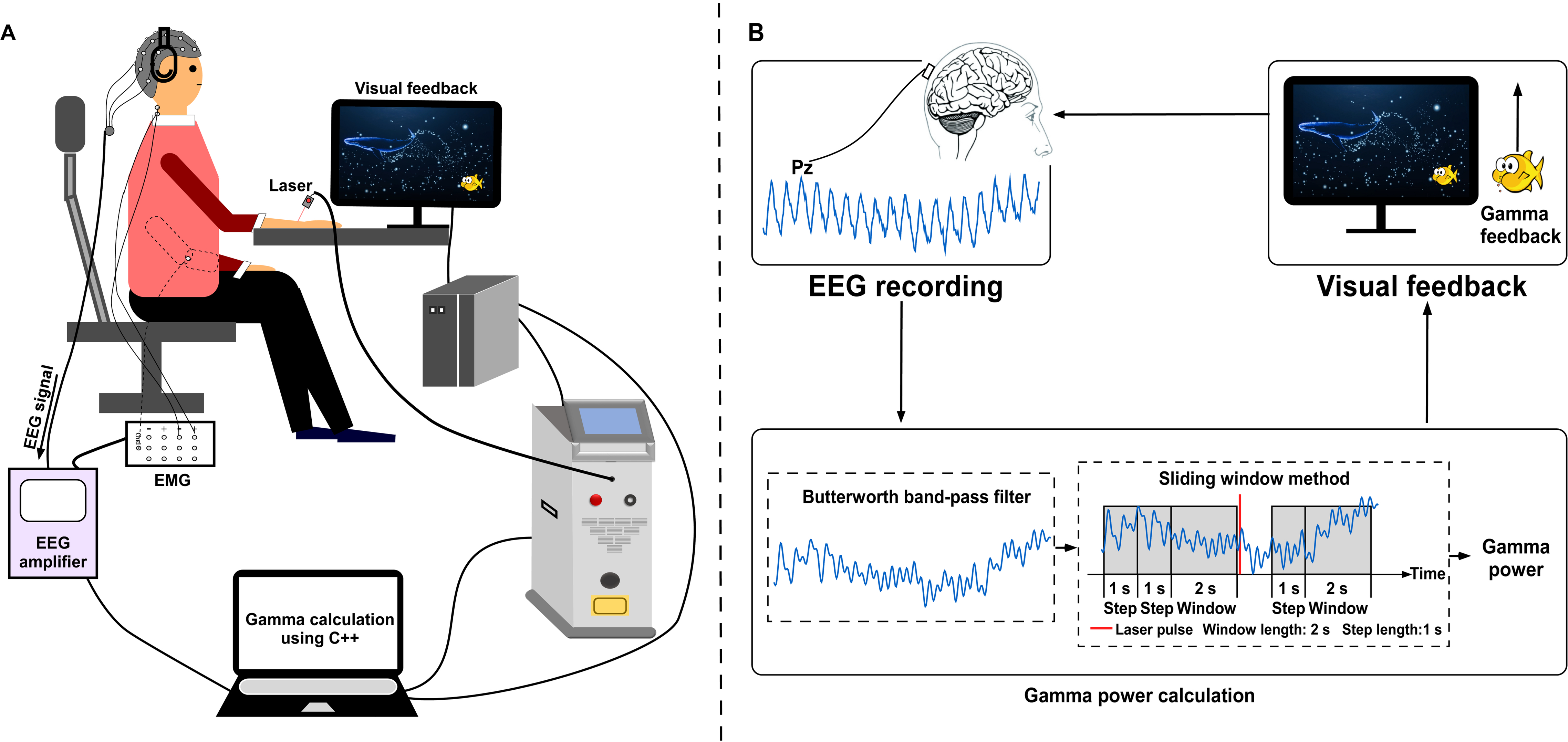
Neurofeedback (NFB) training system. (A) Experimental setup. The participant sat comfortably in a chair, wearing headphones, and maintained a fixed gaze on a computer monitor positioned 30 cm away while watching a video. Nociceptive laser stimuli were delivered to the dorsum of the participant’s right hand. Throughout the entire NFB training, real-time EEG and EMG (in the second part of the experiment) signals were recorded, processed, and used to provide feedback to the participant. (B) Technical implementation diagram. The system consisted of three modules: EEG recording, gamma power calculation, and visual feedback.

##### Module 1: EEG and EMG recording

Subjects were seated in a comfortable chair in a silent, temperature-controlled room. EEG data were recorded using 32 Ag/AgCl electrodes placed according to the international 10-20 system (ANT Neuro, the Netherlands), with a bandpass filter of 0.01-100 Hz and a sampling rate of 2000 Hz. The FCz electrode served as the online reference, and all electrode impedances were maintained below 10kΩ. Real-time EEG signals from the Pz electrode (referenced to the average of M1 and M2) were collected via an EEG amplifier using the Software Development Kit (SDK) protocol. To exclude the possibility that the gamma activity observed over the parieto-occipital region reflected EMG contamination rather than neural activity, an additional electrode was placed over the posterior neck to record EMG activity during the second part of the experiment.

##### Module 2: gamma power calculation

This module was developed in C++ using the BrainAmp SDK. To remove power line interference and isolate gamma oscillations, raw EEG signals from the Pz electrode were first notch filtered between 48-52 Hz and then band-pass filtered between 30-60 Hz using a unidirectional second-order causal Butterworth filter applied online. Prior to initiating NFB, a 10 s baseline of gamma power was estimated by measuring the average power during the initial 10 s of video watching (i.e., the squared amplitudes per second, detailed below). Once the baseline was determined, the NFB began, and a 2-s sliding window was used to estimate real-time gamma power, and this calculation was updated every second to provide immediate feedback. To avoid the influence of stimulus-evoked gamma oscillations, gamma power was not calculated for the 1-s period following each laser stimulation.

##### Module 3: visual feedback

During NFB training, subjects received real-time gamma power feedback through the vertical positioning of an animated fish (in the first scene) or bird (in the second scene) on the displayed video. Specifically, subjects wore headphones and maintained a fixed gaze at a computer monitor placed ∼30 cm away to display videos. The video presentation lasted ∼16 min in total and consisted of two distinct ∼8 min scenes. The first scene depicted a deep ocean environment featuring whales, tridacna, and other rare marine organisms, accompanied by gentle nature sounds. The second scene showcased a seaside environment with flying seagulls, sailboats, and rhythmic waves, along with relaxation instructions for abdominal breathing and muscle relaxation. In the deep ocean scene, the position of the animated fish served as the visual feedback of gamma power changes, while in the seaside scene, a bird served this function. Subjects were instructed to self-regulate their gamma power by focusing their attention and immersing themselves more deeply in the experience, thereby attempting to elevate the animated fish or bird on the screen. No additional guidelines on regulation strategies were provided.

The real-time visual feedback (i.e., the position of the animated fish or bird) is dynamically adjusted based on the difference between current gamma power and the baseline. This difference was continuously transmitted from the measurement equipment to the control computer via socket communication. Positive differences caused the feedback indicator (i.e., a fish or bird) to move upward, while negative differences caused it to move downward. The extent of the movement was proportional to the difference value of gamma power. This dynamic feedback enables subjects to perceive and regulate their gamma power in real-time, with the goal of maximizing it.

### 2.3 Offline EEG and EMG data processing

#### 2.3.1 EEG and EMG data preprocessing

EEG data were preprocessed using EEGLAB^44^. Continuous EEG data were bandpass filtered between 1 and 100 Hz. A notch filter was performed between 48 and 52 Hz to remove power line interference. For spectral analysis, EEG epochs were extracted using a window analysis time of 8000 ms before the stimulus onset (from −8000 to 0 ms). EMG data were preprocessed using the same procedures and parameters as for the EEG data. To extract laser-evoked potentials (LEPs), EEG epochs were extracted using a window analysis time of 3000 ms (from −1000 to 2000 ms) and baseline corrected using the pre-stimulus interval. Trials contaminated by eye blinks and movements were corrected using independent component analysis (runica). After preprocessing, EEG epochs were re-referenced to the average of bilateral mastoids (M1 and M2).

#### 2.3.2 Spectral analysis

An offline spectral analysis was conducted to verify the effect of NFB on *spontaneous* brain oscillations, with a particular emphasis on gamma frequencies. Preprocessed EEG data (from −8000 to 0 ms relative to stimulus onset) were transformed into the frequency domain using a discrete Fourier transform. This procedure yielded EEG power spectra density (PSD) ranging from 1 to 100 Hz for each NFB training session and each subject. These individual PSD were then averaged across subjects within each group, producing session-level spectra of *spontaneous* brain oscillations. To quantify *spontaneous* gamma oscillations, the mean spectral power at Pz electrode within the 30-60 Hz range was extracted. EMG spectral power was quantified using the same procedures and parameters as those used to extract gamma power from the EEG data.

#### 2.3.3 LEP extraction

Preprocessed EEG data were further lowpass filtered at 30 Hz to remove high-frequency components. Single-subject LEP waveforms were obtained by averaging all trials for each subject and each NFB training session. The N2 and P2 waves were identified as the most negative and positive deflections occurring between 0 and 500 ms after stimulus onset, respectively. N2 and P2 amplitudes were measured as the average voltage within a 25-ms window centered on their respective peaks at the Cz electrode. To eliminate the potential influence of linear drift on amplitude estimates, the N2-P2 peak-to-peak amplitude was also calculated for each subject.

### 2.4 Statistical analysis

To evaluate the modulation of NFB training on subjective pain ratings and neural activities across three training sessions, a series of two-way mixed-design analyses of variance (ANOVA) were performed, with a between-subject factor (“group”: active and sham NFB) and a within-subject factor (“session”: session 1, session 2, and session 3). Post hoc tests were performed following any significant main effect or interaction, with false discovery rate (FDR) applied for multiple comparisons (*p* < 0.05). To determine whether the NFB training produced lasting analgesic effects, another two-way mixed-design ANOVA was performed on pain intensity and unpleasantness, with a between-subject factor (“group”: active and sham NFB) and a within-subject factor (“time”: before and after training). Post hoc tests were conducted whenever there was a significant main effect or interaction, with FDR applied for multiple comparisons (*p* < 0.05). Pearson correlation analyses were carried out to explore the relationship between *spontaneous* gamma power and subjective pain ratings in each NFB training session (Bonferroni correction, *p* < 0.05/3 = 0.017). Moreover, Pearson correlation analyses were performed between gamma power at the Pz electrode and gamma power estimated from the EMG electrode, separately for the active and sham groups and for each training session.

All statistical analyses were performed using SPSS 22 (IBM Corp, Armonk, NY, USA), and statistical significance was set at *p* < 0.05.

## 3. Results

### 3.1 The effect of NFB training on spontaneous gamma oscillations

The active and sham groups showed comparable ratings of motivation (active: 5.56 ± 0.96, sham: 4.89 ± 1.29; *p* = 0.538), effort (active: 5.67 ± 1.33, sham: 5.33 ± 1.33; *p* = 0.727), and perceived feedback authenticity (active: 4.44 ± 1.26, sham: 4.33 ± 1.76; *p* = 0.262), suggesting similar task engagement and no obvious between-group differences in the perceived credibility of the feedback. In addition, the response rate in the active group was approximately 52% (23 out of 44), defined as the proportion of subjects who exhibited an increase in gamma power after neurofeedback training. These 23 subjects were then matched, in chronological order, with 23 temporally consecutive subjects from the sham NFB group.

The PSD of *spontaneous* brain oscillations for both groups across the three NFB training sessions is displayed in Figure 3A. Two-way mixed-design ANOVA revealed a significant “session” × “group” interaction for *spontaneous* gamma power (30-60Hz), primarily in the parieto-occipital region (F_(2,86)_ = 6.936, *p* = 0.004, η² = 0.136; Figure 3B). Post hoc paired-sample t-tests showed no significant between-group differences (active vs. sham) at any session (*p* > 0.05). However, in the active NFB group, *spontaneous* gamma power was significantly increased after the third training session compared with both the first (*p* < 0.001) and second sessions (*p* = 0.017). These results suggest that three sessions of NFB training can effectively enhance *spontaneous* gamma oscillations in the parieto-occipital region. Crucially, no significant correlations were observed between gamma power at Pz and that recorded from the EMG electrode in both groups across sessions (Supplementary Figure 1), indicating that the observed changes of gamma activity in the parieto-occipital region are unlikely to be explained by muscle activity.

**Figure 3.**
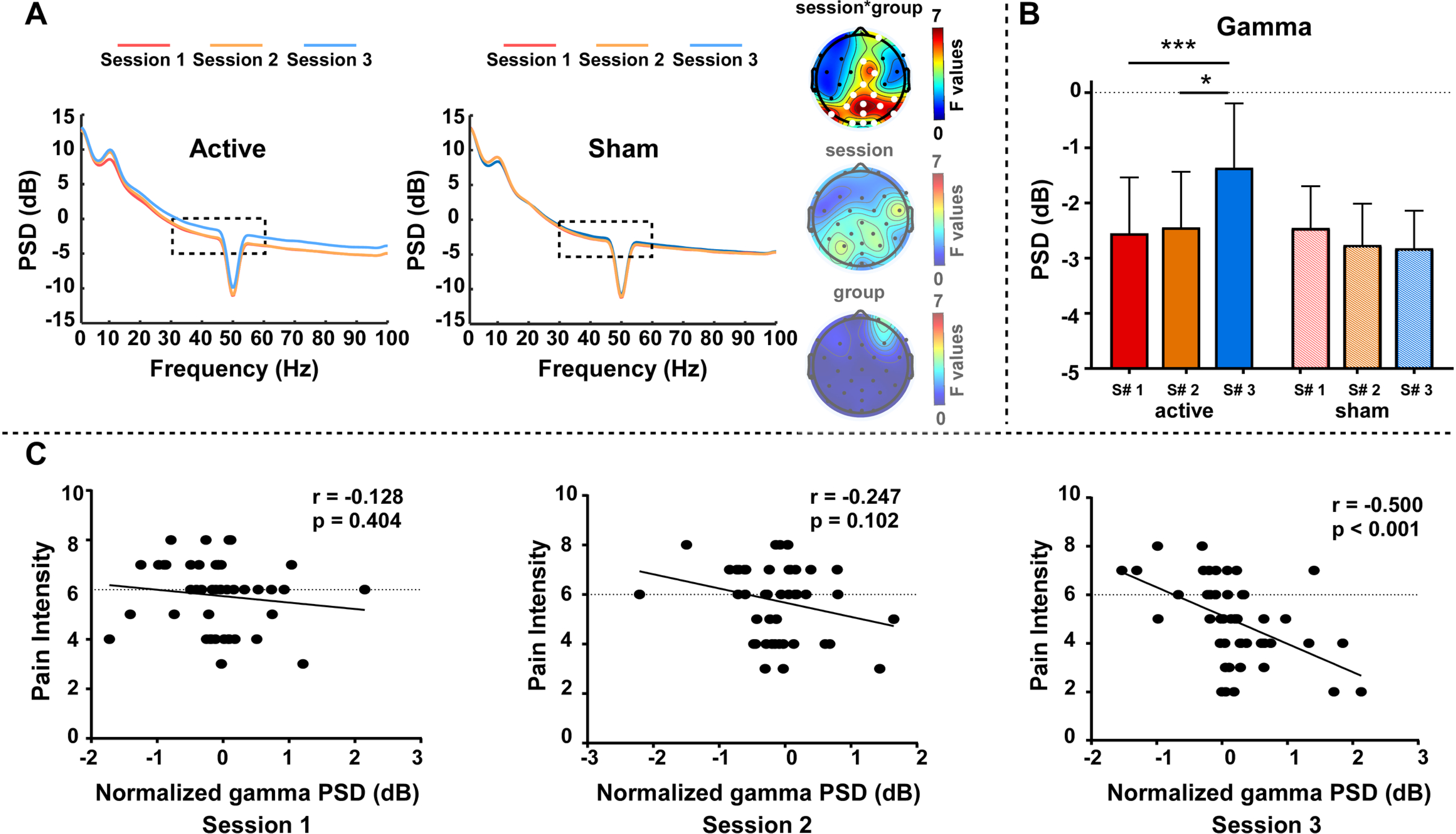
Spontaneous gamma oscillation changes during neurofeedback (NFB) training. (A) PSD of spontaneous EEG oscillations during three NFB training sessions for both active (left) and sham (right) groups, along with scalp topographies of statistical F values for gamma oscillations (30-60Hz). Electrodes with significant effects are marked with white dots (FDR correction, *p* < 0.05). (B) Comparisons of spectral powers of spontaneous gamma oscillations across three NFB training sessions for both groups. (C) Correlations between spectral powers of spontaneous gamma oscillations and pain intensity ratings in three NFB training sessions for the active group. Error bars represent standard errors. *: *p* < 0.05; ***: *p* < 0.001

### 3.2 The effect of NFB training on subjective pain ratings

Two-way mixed-design ANOVA revealed a significant main effect of “session” on both pain intensity (F_(2,86)_ = 9.685, *p* < 0.001, η² = 0.311) and unpleasantness (F_(2,86)_ = 5.427, *p* = 0.008, η² = 0.202). Moreover, a significant “session” × “group” interaction was observed for both ratings (Intensity: F_(2,86)_ = 12.948, *p* < 0.001, η² = 0.376; Unpleasantness: F_(2,86)_ = 4.970, *p* = 0.011, η² = 0.188; Figure 4A). Post hoc paired-sample t-tests showed that, in the active NFB group, both pain intensity and unpleasantness were significantly reduced after the third training session compared with the first (Intensity: *p* < 0.001; Unpleasantness: *p* = 0.001) and second sessions (Intensity: *p* < 0.001; Unpleasantness: *p* = 0.001). In contrast, no significant changes were observed across sessions in the sham NFB group. Notably, after the third training session, the active group reported significantly lower pain intensity than the sham group (*p* < 0.001). These findings suggest that three sessions of NFB training can effectively reduce subjective pain intensity and unpleasantness. Notably, the PSD of *spontaneous* gamma oscillations prior to nociceptive stimuli was significantly correlated with subjective pain intensity in the active group after the third training session (*r* = −0.500, *p* < 0.001; Figure 3C).

**Figure 4.**
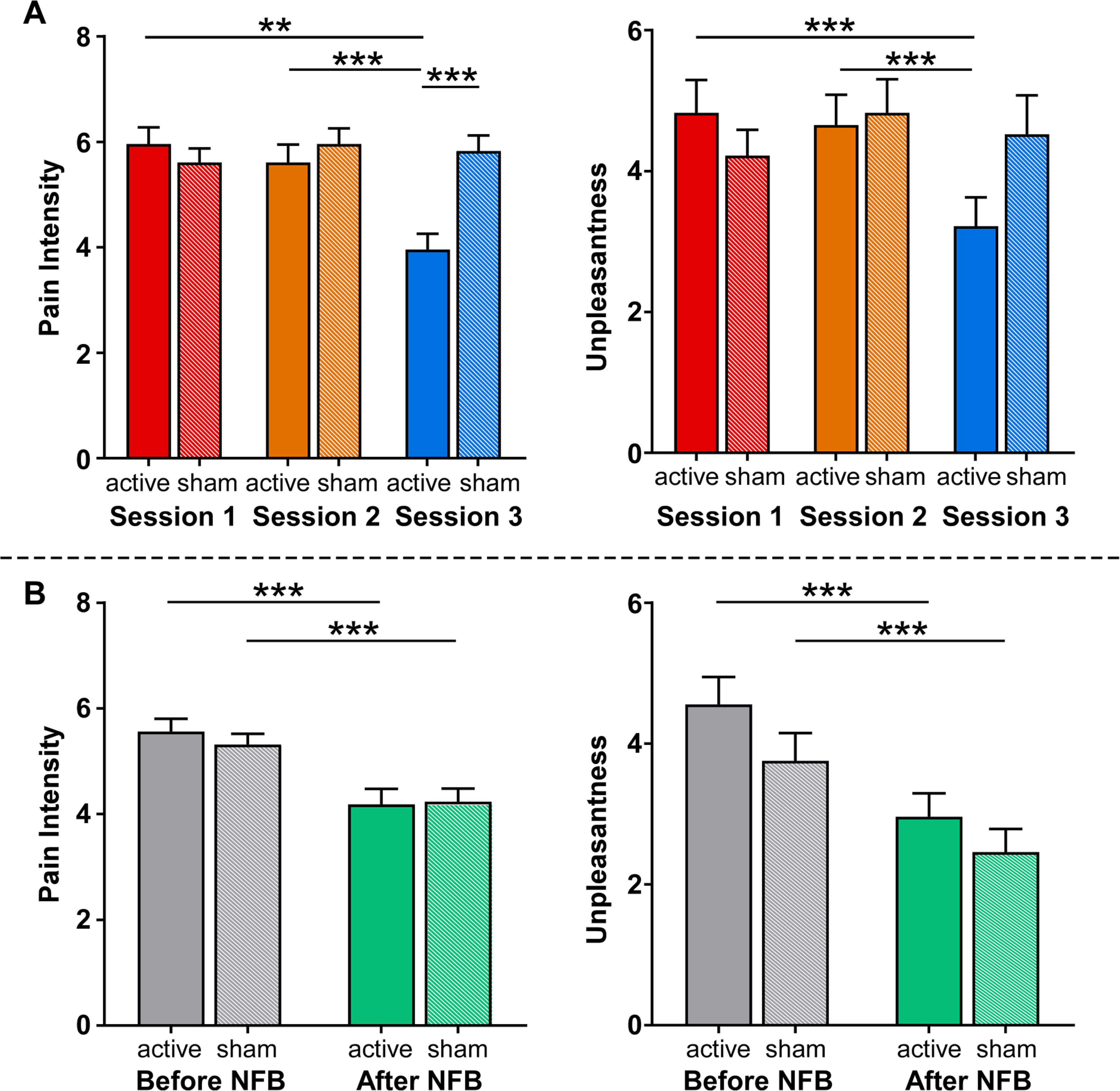
Subjective pain intensity and unpleasantness changes induced by neurofeedback (NFB) training. (A) Comparisons of subjective ratings of pain intensity (left) and unpleasantness (right) across three NFB training sessions for both groups. (B) Comparisons of subjective ratings of pain intensity (left) and unpleasantness (right) before and after NFB training for both groups. Error bars represent standard errors. *: *p* < 0.05; **: *p* < 0.01; ***: *p* < 0.001.

When assessing the lasting effects of NFB training, a significant main effect of “time” was observed for both pain intensity (F_(1,43)_ = 76.164, *p* < 0.001, η² = 0.634) and unpleasantness (F_(1,43)_ = 64.810, *p* < 0.001, η² = 0.596; Figure 4B). Post hoc paired-sample t-tests indicated that both groups reported significantly lower pain intensity and unpleasantness after NFB training (all *p* < 0.001), possibly reflecting the effect of video presentation or an end-of-session effect. However, no significant between-group differences were observed for pain intensity and unpleasantness either before (*p* = 0.446 and *p* = 0.154, respectively) or after NFB training (*p* = 0.898 and *p* = 0.289, respectively). These findings suggest that, although three NFB training sessions reduced pain ratings, the effect was transient, as no lasting group differences were evident at follow-up.

### 3.3 The effect of NFB training on LEP responses

Group-level LEP responses and scalp topographies of N2 and P2 waves for both groups across the three NFB training sessions are presented in Figure 5A. The N2 wave exhibited maximal activity at the vertex, extending bilaterally toward the temporal regions, and the P2 wave was more centrally distributed. Two-way mixed-design ANOVA revealed a significant main effect of “session” for P2 amplitude (F_(2,86)_ = 3.494, *p* = 0.035, η² = 0.077). Additionally, a significant “session” × “group” interaction was observed for N2 amplitude (F_(2,86)_ = 4.261, *p* = 0.025, η² = 0.088) and N2-P2 peak-to-peak amplitude (F_(2,86)_ = 4.619, *p* = 0.019, η² = 0.095). Post hoc paired-sample t-tests indicated that, in the active NFB group, N2 amplitude was significantly reduced after the third training session compared with the first (*p* = 0.007) and second sessions (*p* = 0.004). In the third session, both N2 (*p* = 0.021) and P2 (*p* ≤ 0.001) amplitudes were significantly lower in the active group than in the sham group. Similarly, N2-P2 peak-to-peak amplitude was significantly reduced after the third training session relative to the first (*p* = 0.019) and second sessions (*p* = 0.007). In the third session, the N2-P2 peak-to-peak amplitude was significantly lower in the active group than in the sham group (*p* = 0.001). The sham group did not exhibit any significant changes across sessions. These results suggest that three NFB training sessions can effectively attenuate laser-evoked brain responses, thereby providing converging evidence for the efficacy of NFB training in modulating pain perception.

**Figure 5.**
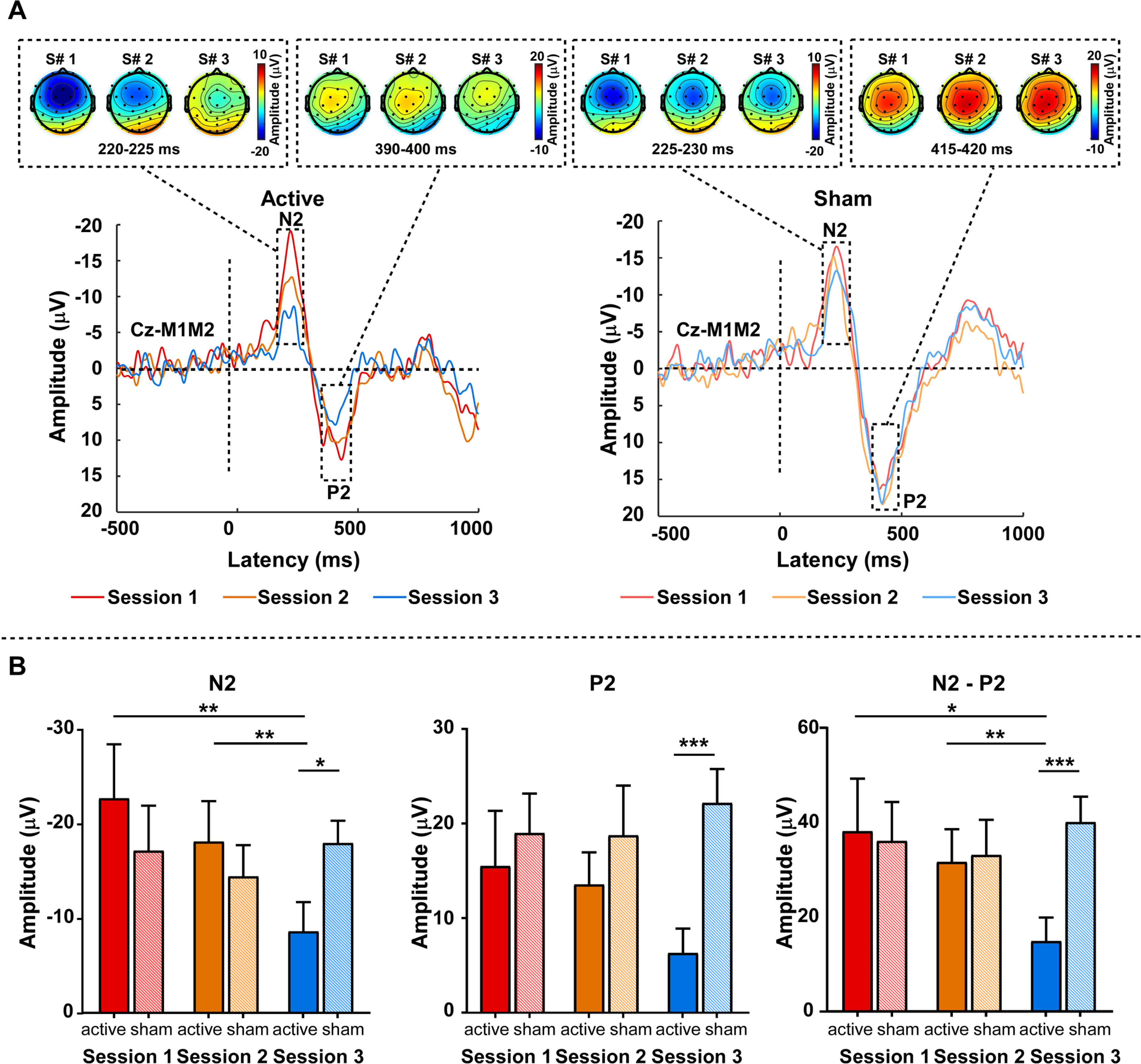
Group-level laser-evoked potentials (LEPs) during neurofeedback (NFB) training. (A) Group-level LEP waveforms at Cz (referenced to bilateral mastoids) and scalp topographies of N2 and P2 waves during three NFB training sessions for both active (left) and sham (right) groups. (B) Comparisons of N2, P2, N2-P2 peak-to-peak amplitudes across three NFB training sessions for both groups. Error bars represent standard errors. *: *p* < 0.05; **: *p* < 0.01; ***: *p* ≤ 0.001.

## 4. Discussion

In this study, we developed a novel NFB training protocol aimed at increasing *spontaneous* gamma oscillations in the parieto-occipital region to alleviate pain. Using a three-session NFB training protocol, we found that approximately 52% of subjects successfully increased *spontaneous* gamma power in the parieto-occipital region. Importantly, the successful modulation of gamma oscillations was associated with a significant reduction in subjective pain ratings, which was further corroborated by decreased laser-evoked brain responses. Theoretically, these findings provided compelling evidence for a causal role of *spontaneous* gamma oscillations in the parieto-occipital region in shaping pain perception. Practically, our study introduced a novel, mechanism-driven NFB therapeutic strategy with potential for effective pain management.

Nociceptive-*evoked* gamma oscillations in the S1 have emerged as a promising pain biomarker, given their ability to track pain intensity selectively regardless of confounding factors such as stimulus saliency^26,29^. Neuromodulation techniques, such as tACS, have therefore been proposed as a potential strategy to modulate gamma oscillations to influence pain perception. However, a study by May et al^37^ that applied tACS at gamma frequency over the somatosensory cortex during tonic experimental pain failed to observe significant analgesic effects. This may be due to the fact that tACS at gamma frequency could not effectively modulate gamma oscillations in the S1 during or immediately before pain perception^37^.

Interestingly, in a large sample of healthy subjects, we observed that prestimulus gamma oscillations in the parietal region were negatively associated with both subjective intensity of pain perception and the brain responses elicited by subsequent nociceptive stimuli^28^. This observation was further supported by a follow-up study, which demonstrated that increased *spontaneous* gamma oscillations prior to contact heat painful stimuli in the parieto-occipital region, induced by an immersive virtual reality scene, were closely linked to subsequent pain relief^36^.

Clearly, *spontaneous* gamma oscillations and nociceptive-*evoked* gamma oscillations are quite different, as they originated from distinct brain regions and exhibited different relationships with pain perception^30^. Instead of targeting nociceptive-*evoked* gamma oscillations in the S1, we directly modulated *spontaneous* gamma oscillations in the parieto-occipital region using a novel NFB strategy. This strategy not only significantly enhanced *spontaneous* gamma oscillations but also reduced pain perception after the third NFB training session. Importantly, *spontaneous* gamma oscillations recorded prior to nociceptive stimuli were significantly and negatively correlated with subjective pain intensity after the third NFB training session. Together, these findings provide evidence supporting a causal relationship between *spontaneous* gamma oscillations in the parieto-occipital region and pain perception.

Unlike nociceptive-*evoked* gamma oscillations in the S1^33^, the increased *spontaneous* gamma oscillations observed in the present study likely reflect a heightened state of cognitive engagement or immersion induced by the NFB training. During the online training, subjects were instructed to self-upregulate their gamma power by focusing their attention and immersing themselves more deeply in the task. This interpretation aligns with prior findings that *spontaneous* gamma oscillations were associated with the sense of immersion induced by a virtual reality scene^36^, emphasizing the effectiveness of engagement and immersion in modulating of *spontaneous* gamma oscillations^45^. Converging evidence from mediation and mindfulness studies further supports this view, with increased parieto-occipital gamma oscillations reported in in experienced practitioners of various meditation practices^46,47^. Crucially, to address the potential confound of muscle activity, the second part of our experiment included simultaneous EMG and EEG recordings. A posterior neck EMG electrode was incorporated because neck muscle activity represents a plausible source of high-frequency contamination in the parieto-occipital region. The results revealed no significant correlations between EEG and EMG gamma powers in either group for all sessions, indicating that the gamma activity observed at Pz is unlikely to be driven by muscle activity. Although we cannot entirely exclude all possible muscle artifacts, this additional experiment provides direct evidence that muscle activity is unlikely to be the primary contributor to the observed gamma power at Pz.

Utilizing real-time EEG monitoring to provide feedback, NFB enables subjects to visualize and directly modulate their brain activity in a targeted manner, thereby facilitating an explicit, observable, self-regulatory process^39^. Consequently, NFB offers distinct advantages over other neuromodulation techniques, such as tACS, by providing personalized, real-time modulation of brain functions based on individuals’ ongoing neural activity patterns^48^. Through repeated training, subjects may acquire the ability to self-regulate these neural signals even in the absence of continuous EEG feedback, potentially supporting sustained neural control without reliance on external stimulation or devices. In this study, our NFB training strategy not only successfully enhanced *spontaneous* gamma oscillations but also reduced subjective pain perception and nociceptive-evoked brain responses, highlighting its effectiveness in pain modulation. Additionally, our results supported the view that repeated NFB training is crucial for improving the modulation efficiency^38^. Specifically, we found that *spontaneous* gamma oscillations, pain perception ratings, and nociceptive-evoked neural responses did not significantly change after the first and second NFB training sessions, whereas evident effects emerged after the third NFB training session. This pattern indicated that robust changes in both *spontaneous* gamma oscillations and pain perception likely required repeated learning and consolidation. Given that neural plasticity requires repeated training^38,49^, increasing the number of training sessions may help consolidate the neural adaptations necessary for sustained modulation of gamma oscillations, thereby enhancing the potential for evident reductions in pain perception^49^. It is also noteworthy that both groups exhibited reduced pain perception ratings after the entire NFB training. Prior research has demonstrated that visual stimuli and attentional engagement can influence pain perception^36,50^. Therefore, the observed changes in pain perception in both groups may reflect nonspecific factors, such as the modulatory effects of video viewing or end-of-experiment effects, rather than NFB-specific mechanisms.

Several limitations should be acknowledged. First, although this study introduces a novel strategy to modulate pain perception, the analgesic effects of NFB were only observed after three sessions of training in healthy subjects and did not persist at follow-up. Further investigation is needed to improve the long-term efficacy of gamma-based NFB. Second, although posterior neck EMG recordings were included to control for potential muscle contamination of gamma-band EEG activity, this approach cannot completely exclude all possible EMG contributions. Nevertheless, the absence of significant correlations between Pz gamma power and posterior neck EMG gamma power suggests that the observed modulation at Pz is unlikely to be driven by muscle activity. Third, the analgesic effects were mainly observed in active NFB responders, defined as subjects who successfully increased gamma activity after training. Therefore, the present findings should not be generalized to all individuals receiving gamma-based NFB. Future studies should identify predictors of NFB responsiveness and develop strategies to reduce the proportion of non-responders. Fourth, while modulating *spontaneous* gamma oscillations in the parieto-occipital region was associated with reduced pain perception, the underlying mechanisms remain unclear, particularly how *spontaneous* gamma oscillations in the targeted brain region influence nociceptive-evoked brain responses. Fifth, the analgesic effects of NFB were evaluated solely in healthy subjects. Further studies are needed to evaluate the clinical utility of this approach in patient populations experiencing chronic or pathological pain.

## Data availability statement

The data have been publicly uploaded and are available at: https://www.scidb.cn/s/YjANBf.

## Authors’ contributions

XYZ: designed and conducted the experiments, analyzed data, and wrote the first draft.

HS, HQD, SYW, GH: assisted with data collection.

JWL, HLW, XJL: assisted with data analyzation.

YZB, LH: designed the experiments, wrote the first draft, edited, and reviewed the manuscript.

## Acknowledgements

This work was supported by the Beijing Natural Science Foundation (5252019, JQ22018) and National Natural Science Foundation of China (32071061).

## Supplementary Materials

**Supplementary Figure 1.** Correlations between gamma power at Pz and that recorded from the EMG electrode. No significant correlations were found in either the active or sham group across sessions, indicating that the modulation of gamma power at Pz is unlikely to reflect muscle activity.

